# Mechanisms of Alpha-Synuclein Seeded Aggregation in Neurons Revealed by Fluorescence Lifetime Imaging

**DOI:** 10.1101/2024.12.16.628520

**Authors:** Paula-Marie E. Ivey, Magaly Guzman Sosa, Abdelrahman Salem, Sehong Min, Wenzhu Qi, Alicia N. Scott, Karin F. K. Ejendal, Tamara L. Kinzer-Ursem, Jean-Christophe Rochet, Kevin J. Webb

**Affiliations:** Weldon School of Biomedical Engineering, Purdue University, 206 S Martin Jischke Dr, West Lafayette, IN 47907, USA; Purdue Institute for Integrative Neuroscience, Purdue University, 207 S Martin Jischke Dr, West Lafayette, IN, 47907, USA; Borch Department of Medicinal Chemistry and Molecular Pharmacology, Purdue University, 575 Stadium mall drive, West Lafayette, IN 47907, USA; Elmore Family School of Electrical and Computer Engineering, Purdue University, 465 Northwestern Ave, West Lafayette, IN 47907, USA

## Abstract

The brains of Parkinson’s disease (PD) patients are characterized by the presence of Lewy body inclusions enriched with fibrillar forms of the presynaptic protein alpha-synuclein (aSyn). Despite related evidence that Lewy pathology spreads across different brain regions as the disease progresses, the underlying mechanism hence the fundamental cause of PD progression is unknown. The propagation of aSyn pathology is thought to potentially occur through the release of aSyn aggregates from diseased neurons, their uptake by neighboring healthy neurons via endocytosis, and subsequent seeding of native aSyn aggregation in the cytosol. A critical aspect of this process is believed to involve the escape of internalized aggregates from the endolysosomal compartment, though direct evidence of this mechanism in cultured neuron models remains lacking. In this study, we utilize a custom-built, time-gated fluorescence lifetime imaging microscope (FLIM) to investigate the progression of seeded aggregation over time in live cortical neurons. By establishing fluorescence lifetime sensitivity to aSyn aggregation level, we are able to monitor the protein’s aggregation state. Through a FLIM analysis of neurons expressing aSyn-mVenus and exposed to aSyn preformed fibrils labeled with the acid-responsive dye pHrodo, we reveal the protein’s aggregation state in both the cytosol and the endolysosomal compartment. The results indicate that aSyn seeds undergo partial disassembly prior to escaping the endocytic pathway, and that this escape is closely linked to the aggregation of cytosolic aSyn. In certain neurons, monomeric aSyn is found to translocate from the cytosol into the endolysosomal compartment, where it apparently forms aggregates in proximity to retained seeds. Additional analyses reveals zones of neuritic aSyn aggregates that overlaps with regions of microtubule disruption. Collectively, these findings enhance our understanding of aSyn pathology propagation in PD and other synucleinopathies, motivate additional experiments along these lines, and offer a path to guide the development of disease-modifying therapies.

## 2 Introduction

Several neurodegenerative diseases are characterized by a spread of protein aggregates throughout the brain^1–3^ and involve the release of a fibrillar aggregate, termed a seed, from a diseased neuron and its subsequent internalization by a healthy neuron via endocytosis. ^4^ This seed then induces the templated aggregation of the monomeric cytosolic protein, thereby propagating self-assembly.^5–7^ A pathological hallmark of Parkinson’s disease (PD) and other synucleinopathy disorders is the presence in surviving neurons of Lewy body inclusions enriched with fibrillar forms of the presynaptic protein alpha-synuclein (aSyn), and evidence suggests that Lewy pathology spreads from the brainstem to the midbrain and eventually into the cortex as the disease progresses.^8–10^ However, the precise molecular mechanisms underlying the propagation of aSyn aggregates in PD remain unclear, reflecting similar gaps in our understanding of other neurodegenerative diseases.^11,12^ A key step in the spread of aSyn pathology is believed to be the escape of internalized aggregates from the endolysosomal compartment into the cytosol, a pathway that competes with their clearance via lysosomal autophagy. ^13–16^ Evidence suggests that a loss of membrane integrity of the endolysosomal compartment leads to fibril escape, enabling the recruitment of monomers and subsequent seeded aggregation of the cytosolic protein. ^14,16^ Previous studies have employed co-localization techniques using fluorescently labeled fibrils and markers of endolysosomal rupture, such as fluorescently tagged Galectin-3, to infer fibril escape in immortalized cell lines.^13^ However, these methods do not allow direct visualization of aggregates within the endolysosomal compartment, nor do they capture the real-time coupling between aggregate escape and cytosolic seeding. Furthermore, the fibril escape mechanism has yet to be studied in cultured neurons.

Investigating the underlying mechanisms of aggregation requires highly sensitive methods to probe the evolution of aggregate states in different subcellular compartments in real time. Although reagents with a high affinity for fibrillar aSyn, such as antibodies ^17–19^ or fluorescent dyes (e.g., thioflavin S),^20,21^ are commonly used to monitor aSyn aggregation, their binding is not always specific, and they are unsuitable for use in live cells, limiting their capacity to provide insights into the dynamic evolution of aggregates. As an alternative, fluorescence lifetime measurements of aSyn-fluorescent protein (FP) fusions provide a powerful tool to monitor aSyn self-assembly in live cells. During aggregate formation, the fluorescence lifetime of the FP fused to aSyn decreases due to self-quenching as fluorophores on multiple aSyn-FP subunits come into close proximity, as illustrated in Fig. 1.^22^ Self-quenching has been used to monitor the spontaneous or seeded aggregation of aSyn fused to the yellow FP variant, mVenus, in cell culture models and in the nematode *C. elegans*.^23–26^

**Figure 1:**
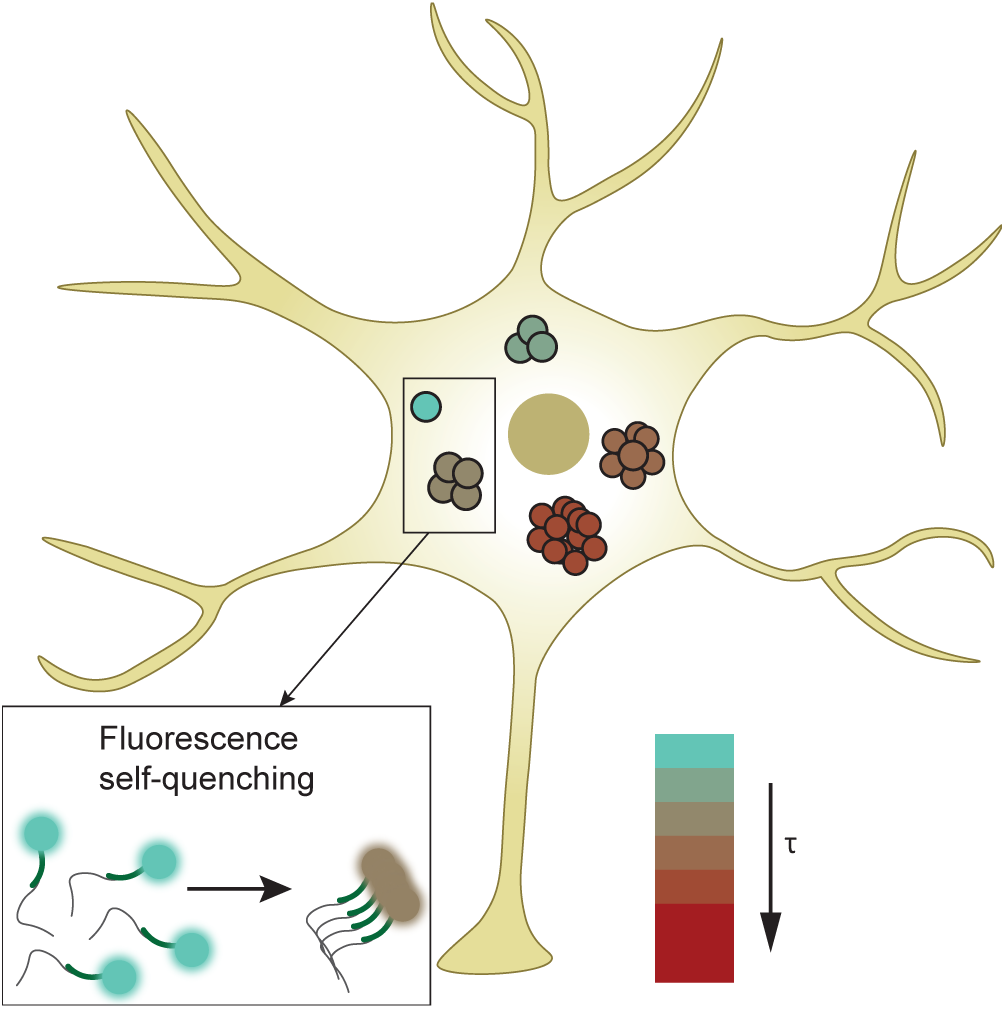
Illustration of the principle of fluorescence lifetime reduction due to aSyn aggregation. As aSyn tagged with a fluorescent dye or protein aggregates, the local concentration of the fluorophore increases, leading to a decrease in fluorescence lifetime through a self-quenching mechanism. This reduction in lifetime is represented by a color gradient shifting from blue (monomeric state) to red (fibrillar state).

In this study, we used fluorescence lifetime measurements to investigate the fate of expressed aSyn-mVenus and internalized fluorescently-labeled seeds in live cortical neurons over several days. This approach enabled us to evaluate the protein’s aggregation state in different subcellular compartments, as well as map zones of aSyn-mVenus aggregation within neuronal processes. Our findings suggest that seeded aSyn aggregation can occur in either the cytosol or the endolysosomal compartment, ultimately leading to the accumulation of neuritic aggregates in areas characterized by microtubule disruption.

## 3 Results and Discussion

### 3.1 Design of Time-Gated FLIM System

A time-gated FLIM system was designed and constructed to carry out fluorescence lifetime measurements in neurons. Figure 2(a) shows the principle, where the fluorescence emission decay curve (red) is incrementally sampled by changing the gate delay of an intensifier relative to the initial laser excitation pulse (green) using an electrical picosecond delay unit. Lifetime information can then be extracted pixel-wise from the sequential set of images collected as a function of temporal gate delay. To carry out the FLIM measurements, a custom time-gated wide-field FLIM system with pulsed laser excitation was constructed around an Olympus IX73 platform (Fig. 2(b), the details of which are described in the methods section).

**Figure 2:**
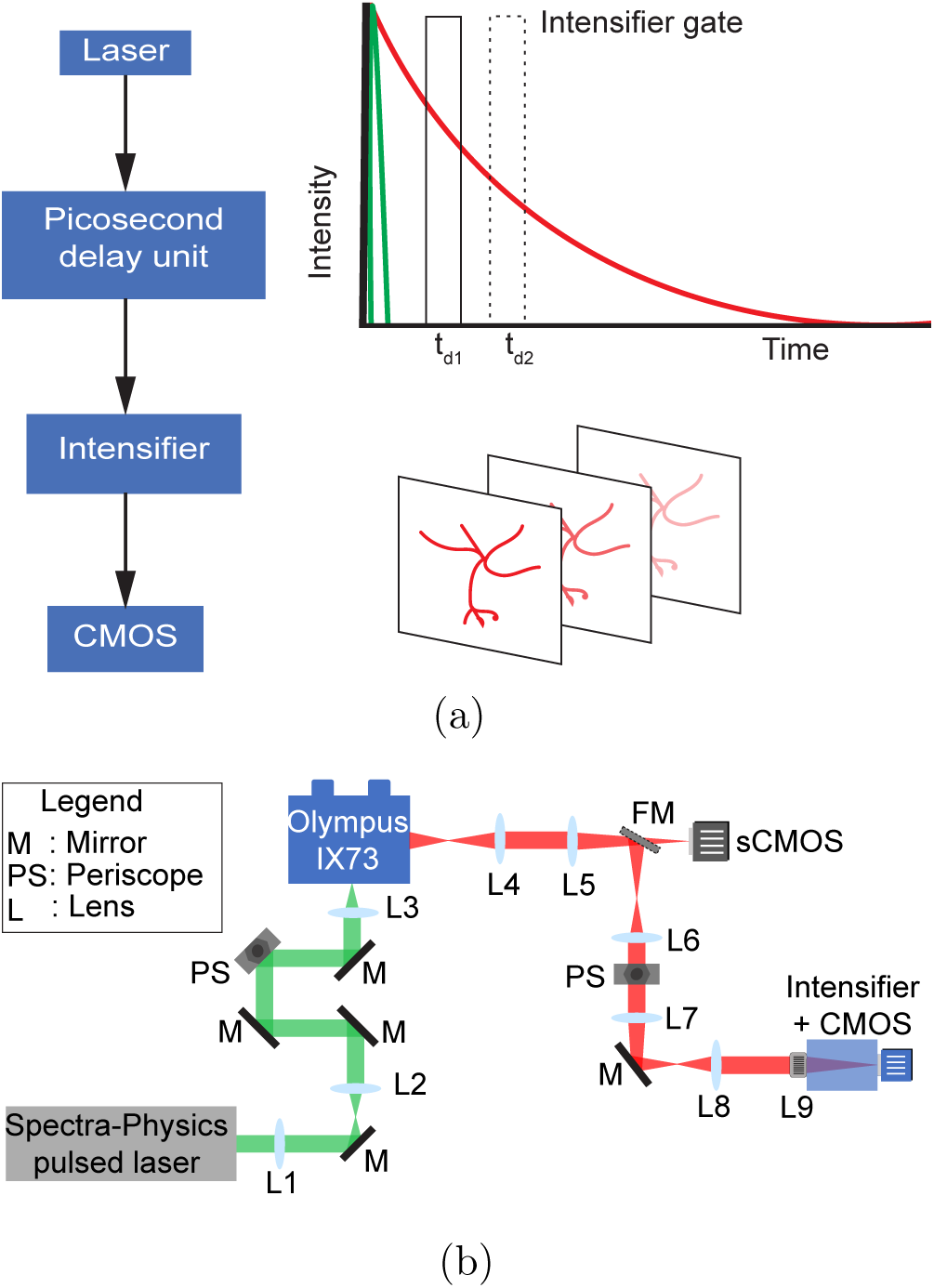
(a) The FLIM measurement principle. An intensifier gate incrementally samples the fluorescence decay curve (red), and a picosecond delay unit changes the temporal gate delay relative to the laser excitation pulse (green). At each gate delay position, an image is generated that is utilized to determine lifetime information in each pixel of the image. (b) Schematic of the time-domain FLIM system used for detecting aggregation (top view, not drawn to scale). A Spectra-physics ultrafast laser and optical parametric amplifier system (Spirit HE 1040-16 pump with OPA 30, yielding approximately 150 fs pulses) provide excitation energy (green) coupled to an Olympus iX73 microscope via a dichroic mirror. The emission path collected from the microscope is depicted in red. L: lens, M: mirror, BS: beam splitter, PS: periscope, FM: flip mirror.

### 3.2 FLIM Analysis Shows Reduced aSyn-mVenus Lifetimes in Fixed Neurons Following PFF Treatment

We first examined the effect of PFF-mediated seeding on the fluorescence lifetime of fluorescently tagged aSyn in fixed neurons. Cortical neurons prepared from the brains of embryonic rats were transduced with adenovirus encoding aSyn A53T-mVenus (hereafter referred to as aSyn-mVenus) under the control of the neuron-specific synapsin promoter. Our rationale for expressing A53T aSyn is that this variant, which is linked to familial PD,^27,28^ undergoes aggregation more rapidly than wild-type aSyn.^29,30^ The transduced neurons were incubated with or without mouse aSyn PFFs for 6 days and then fixed with 4% (w/v) paraformaldehyde (PFA) in the absence or presence of 1% (v/v) Triton X-100, a detergent used to remove soluble protein, leaving only insoluble aggregates.

When imaging fixed PFF-treated neurons, we found the lifetime to be generally shorter than for the untreated samples, indicative of aggregate-driven quenching. Figure 3(a) shows a representative lifetime map for neurons having no PFF treatment and Fig. 3(b) a case for PFF-treated neurons. We note that, as the representative data in Fig. 3(b) indicates (lower left, as opposed to the remainder of this image), there were instances where some neurons treated with PFFs did not have reduced lifetime. Figure 3(c) presents a representative lifetime map, indicating the presence of detergent-resistant aSyn-mVenus aggregates, whereas minimal fluorescence was detected in detergent-fixed cells incubated without PFFs. Through multiple measurements, we found that both the control and PFF-treated samples exhibited a range of lifetimes. However, the reduction in the mean lifetime in neurons cultured with PFFs was found to be statistically significant, compared to those without PFFs, in both detergent-treated and untreated control samples, as is clear in the data in Fig. 3(d). This decrease in mean fluorescence lifetime aligns with findings from a study on PFF-treated HEK biosensor cells expressing aSyn-YFP,^26^ although our results represent the first reported use of FLIM to monitor intracellular seeded aSyn aggregation in a primary neuron model. ^26^

**Figure 3:**
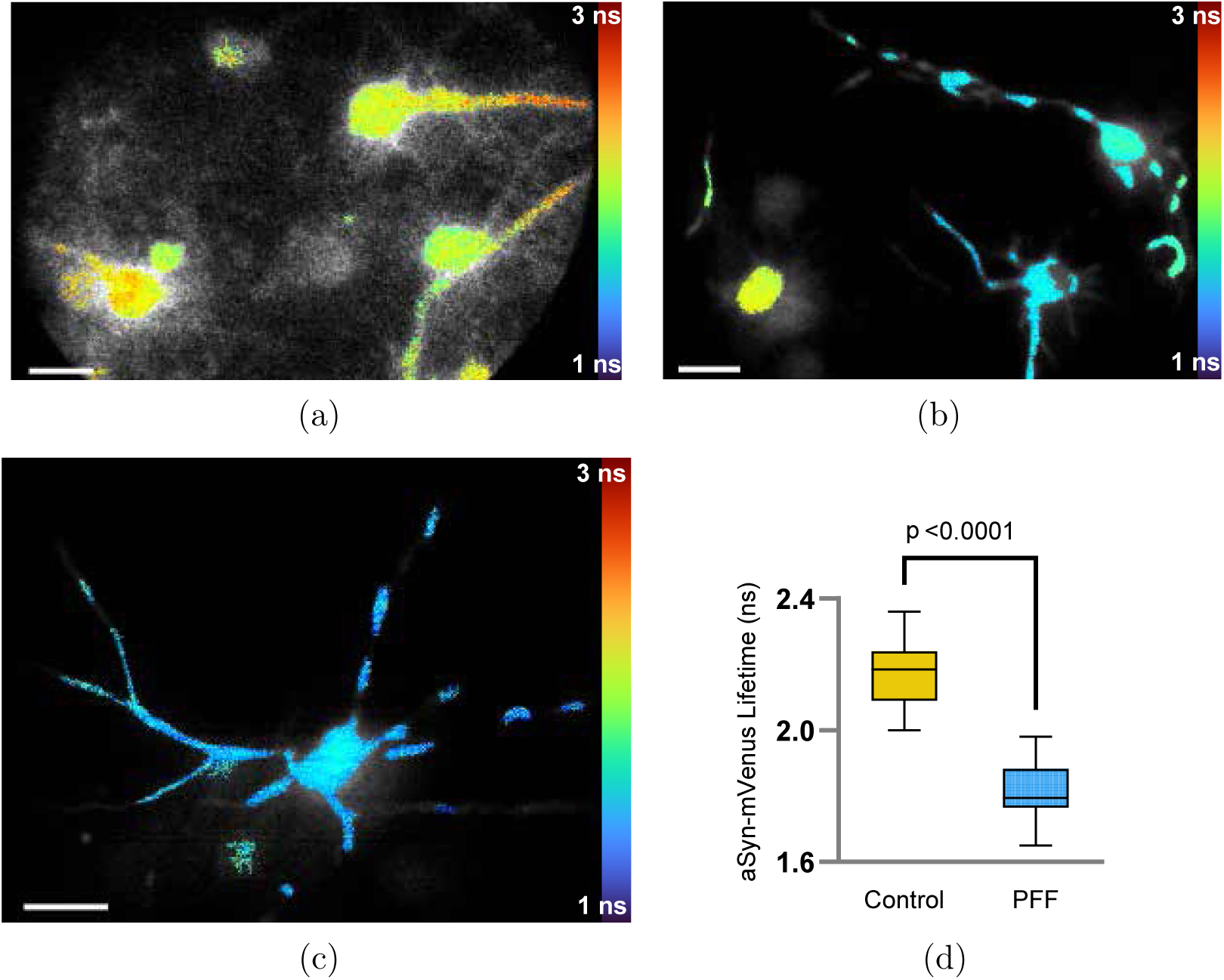
A decrease in aSyn-mVenus lifetime is observed in fixed neurons following PFF treatment. (a)-(c) Representative lifetime maps for rat cortical neurons expressing aSyn-mVenus, fixed and imaged 6 days after incubation without (a) or with (b and c) aSyn PFFs. The neurons were fixed without (a and b) or with (c) Triton X-100 (1%, v/v). Scale bars in panels (a)-(c): 20 *µ*m. (d) Graph showing the fluorescence lifetime for the control versus PFF-treated neurons, fixed without exposure to 1% (v/v) Triton X-100 (*n* = 22 neurons from 3 independent cultures per group), indicating a significant reduction in lifetime for the PFF-treated samples. The data are shown as box plots (*p <* 0.0001, unpaired t-test).

In the lifetime map shown in Fig. 3(b), neurons in the PFF-treated sample are shown with distinct lifetime values, demonstrating the capability of lifetime measurements to differentiate neurons with or without aggregates within the same field of view. These results indicate that aSyn-mVenus was induced to form aggregates by seeds internalized in neurons exhibiting low fluorescence lifetimes, whereas neurons with high lifetime values presumably did not internalize the seeds and thus maintained a predominantly soluble pool of aSyn. Consistent with this idea, PFF-treated cultures fixed with Triton X-100 only displayed low lifetime values. In addition, neurons exhibiting signs of seeded aggregation showed a redistribution of aSyn-mVenus along the axon, producing a dashed intensity pattern that was apparent in samples fixed with or without Triton X-100 (Figs. 3(b) and (c)). This distinct pattern provided a reliable marker for monitoring seeded aSyn-mVenus aggregation in subsequent FLIM studies of live neurons.

### 3.3 FLIM Analysis Reveals a Gradual Decrease in aSyn-mVenus Lifetime in Live Neurons Exposed to PFFs

Having confirmed that FLIM provides a sensitive measure of intracellular seeded aggregation in neurons expressing aSyn-mVenus, we applied this method to monitor aSyn aggregation at various times post-seed administration in live neurons. By integrating a live-cell chamber into our imaging setup, we were able to conduct FLIM measurements on the same neurons every 24 h over a 5-day period, starting 3 days after PFF administration (“Day 1”). Each day, neurons were selected by applying a threshold, and a common thresholded mask was used to process the intensity images for that day. Detailed procedures, including the MATLAB-generated GUI for live-cell image analysis, are presented in Supplementary Fig. 1.

Figure 4 presents the results from this live-cell study. Over the 5-day period, we observed a gradual decrease in fluorescence lifetime in PFF-treated neurons that eventually exhibited aSyn-mVenus pathology on Days 4 and 5, as evidenced by the appearance of a dashed intensity pattern (Figs. 4(a) and (b)), similar to that observed in fixed cells (Figs. 3(b) and (c)). These results underscore the sensitivity of the FLIM method, enabling aggregate detection as early as 4 days after PFF administration (Day 2 in Fig. 4(b)).. A comparison of the mean lifetime distributions (calculated by plotting the number of pixels within specific lifetime bins) across multiple neurons in each group revealed a progressive shift toward lower lifetime values in PFF-treated neurons from Day 1 to Day 5, whereas the lifetime distribution for control neurons remained largely unchanged over the same period (Figs. 4(b) and (c)).

**Figure 4:**
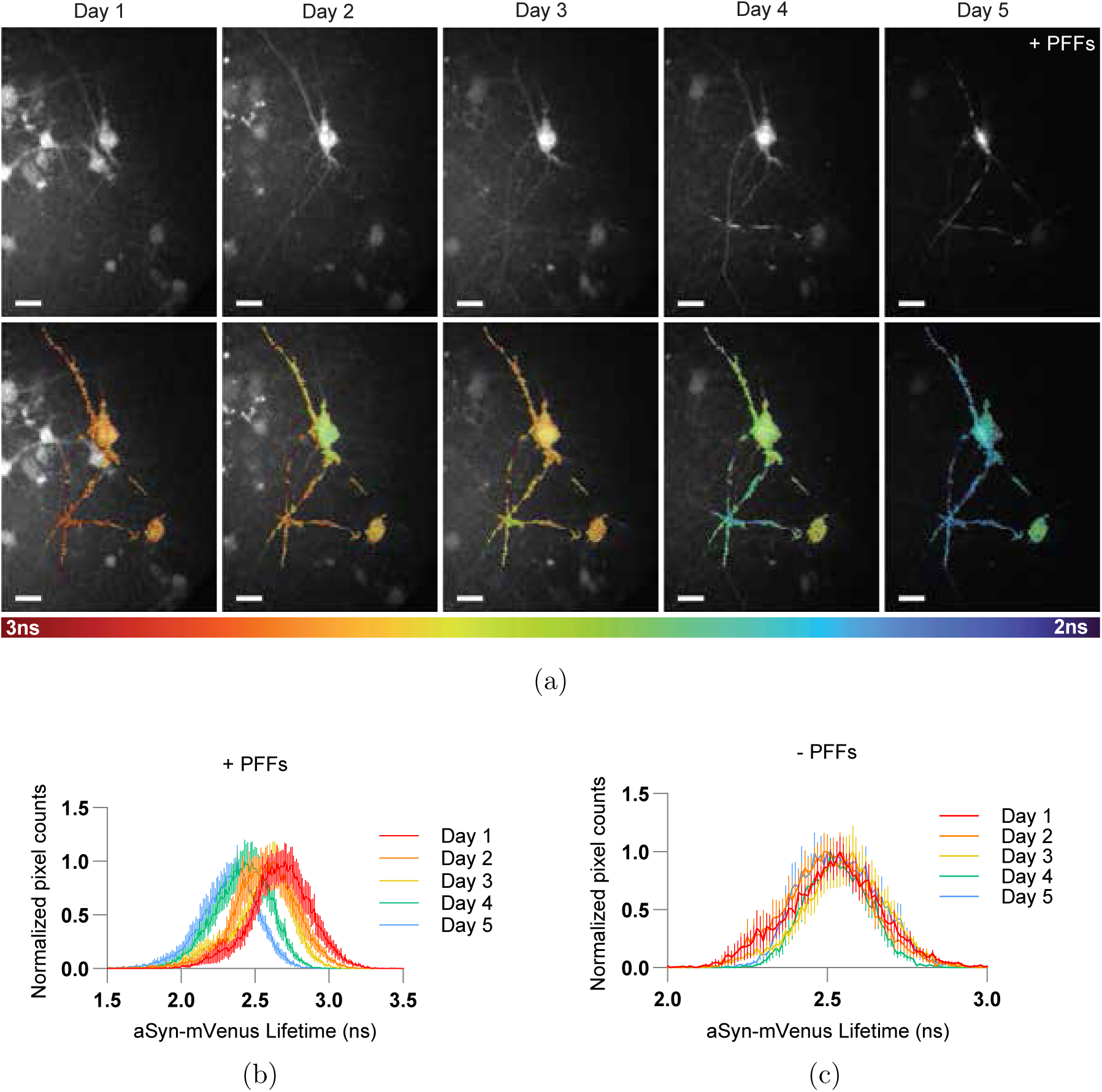
A gradual decrease in aSyn-mVenus lifetime is observed in live neurons exposed to PFFs. (a) Intensity images (top) and lifetime maps (bottom) of representative live cortical neurons expressing aSyn-mVenus, recorded from 3 to 7 days post-PFF treatment (identified as Day 1 through Day 5). In the intensity images (top), two neurons display aggregate formation over time, evidenced by the dashed intensity structures along the axons. Corresponding lifetime maps (bottom) reveal a concomitant decrease in lifetime with aggregate formation. Scale bars: 20 *µ*m. (b, c) Lifetime histograms over 5 days for neurons cultured in the presence (b) or absence (c) of aSyn PFFs. The histograms show lifetime distributions across all pixels contained within neurons (20 neurons from 2 independent cultures). The data are presented as the mean +*/* SEM (SEM: standard error in the mean, 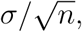 with *σ* equal to the standard deviation for each sample and *n* = 20).

### 3.4 Tracking the Fate of Internalized aSyn Seeds and Cytosolic aSyn-mVenus Reveals Diverse Endolysosomal Mechanisms Underlying the Spread of aSyn Pathology

We now describe the results from a set of FLIM experiments to investigate the connection between the fate of aSyn seeds within the endolysosomal compartment and the efficiency of aSyn-mVenus aggregation in the cytosol, as presented in Fig. 5. To monitor internalized seeds, aSyn PFFs were labeled with pHrodo Red (hereafter pHrodo), a dye that exhibits significantly higher fluorescence in acidic environments. To simulate the expected increase in pHrodo quantum yield in the endolysosomal compartment compared to the cytosol, we measured the dye’s fluorescence intensity at pH 5.5 versus pH 7.4 and found that the intensity was approximately 4-fold greater at the lower pH. We then administered pHrodo-labeled PFFs to rat cortical neurons expressing aSyn-mVenus and imaged the live neurons 6 days post-treatment. Some neurons were found to retain PFFs in the endolysosomal compartment, as indicated by the presence of cellular entities that contained pHrodo-labeled PFFs (top panel of Fig. 5(a), Retention). In contrast, other neurons displayed no pHrodo fluorescence but exhibited a dashed/non-homogeneous aSyn-mVenus fluorescence intensity pattern and reduced aSyn-mVenus fluorescence lifetimes, shorter than those observed in neurons with retained PFFs (Fig. 5(a) bottom panels, Escape (i) and Escape (ii)). The retention cases did not exhibit the dashed fluorescence intensity pattern we found to be characteristic of aSyn-mVenus aggregation in neuronal processes, and the fluorescence lifetime of aSyn-mVenus in these neurons was similar to that observed in control neurons and notably longer than the lifetimes observed in neurons treated with PFFs (compare Fig. 5(a), top panel, Retention and Fig. 5(b) to Figs. 4(a) and (b)). The lifetime statistics in Fig. 5(b) show the substantially reduced aSyn-mVenus lifetime for neurons with escaped aggregates. These results suggest that PFFs were initially internalized in these cells but later translocated from the endolysosomal compartment to the cytosol, where they induced the aggregation of cy-tosolic aSyn-mVenus. The fact that the mean aSyn-mVenus lifetime was significantly lower in neurons with escaped versus retained PFFs (Fig. 5(b)) supports the idea that PFF escape from the endolysosomal compartment is an essential step for inducing aggregation and for the propagation of aSyn aggregates. Our findings are consistent with the results of earlier studies suggesting that the endocytic escape of aSyn seeds, facilitated by the disruption of the endolysosomal membrane, plays a key role in the spread of aSyn pathology in cultured cells.13,14,16,31

**Figure 5:**
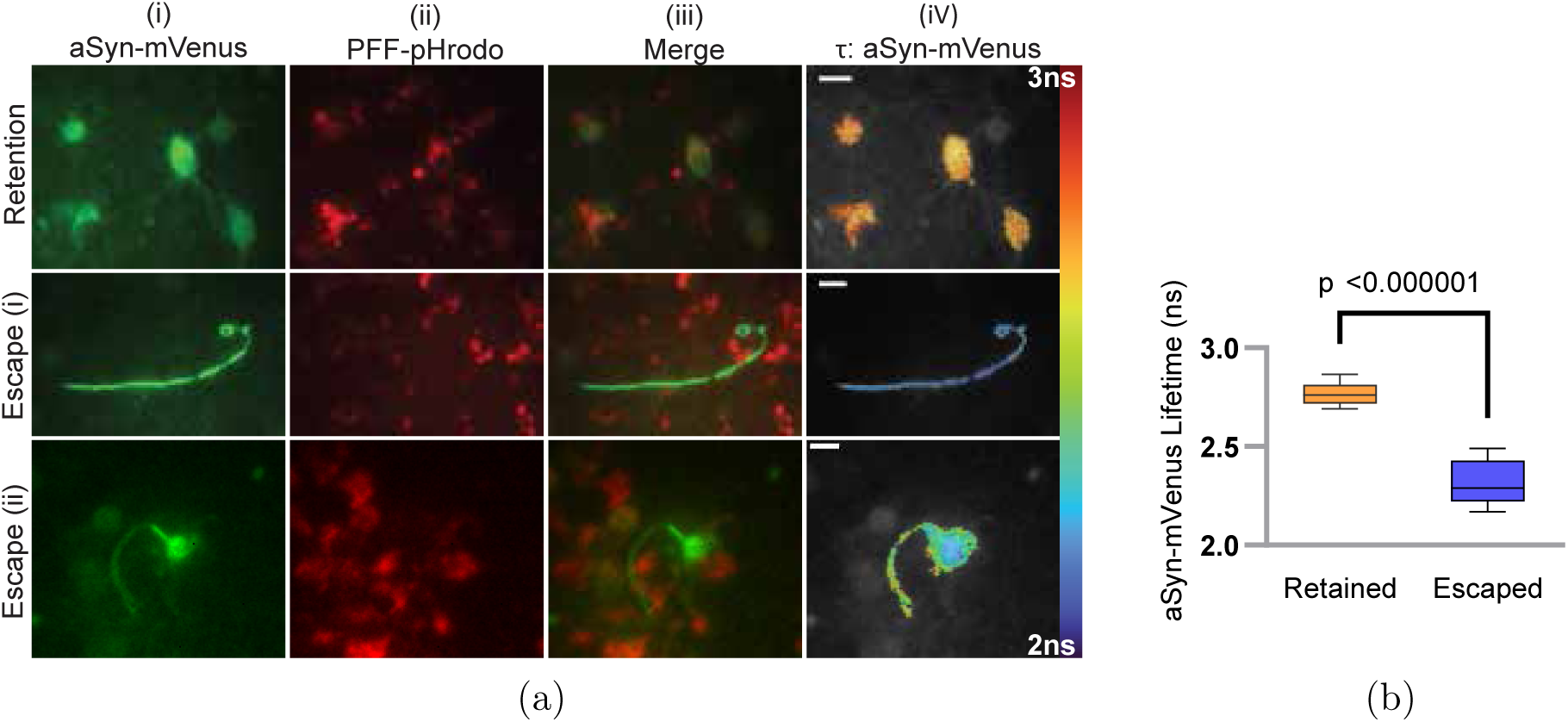
PFF escape from the endolysosomal compartment precedes seeded aggregation. (a) Images of representative live cortical neurons transduced with adenovirus encoding aSyn-mVenus and treated with pHrodo-labeled aSyn PFFs, recorded 6 days post-PFF treatment. The images display the fluorescence intensity of aSyn-mVenus (i) and pHrodo (ii), the merged intensity signals (iii), and the corresponding aSyn-mVenus lifetime map (iv) of neurons showing evidence of seed retention (top row) or escape (bottom rows, Escape (i) and (ii)). Scale bar: 10 *µ*m. (b) Graph showing the aSyn-mVenus fluorescence lifetime for neurons with retained or escaped PFFs (*n* = 12 neurons), indicating a significant reduction in lifetime for neurons with escaped seeds. The data are shown as box plots (*p <* 0.000001, unpaired t-test).

Additional FLIM studies were conducted to investigate the interplay between seed escape/retention and aSyn-mVenus aggregation over time. Rat cortical neurons expressing aSyn-mVenus were treated with pHrodo-labeled PFFs and imaged every 24 h for 4 days, starting on Day 1, which corresponded to 3 days post-PFF administration. These experi-ments were performed using two independent cultures: one treated with human A53T aSyn PFFs and the other with mouse aSyn PFFs.

In general, neurons treated with human aSyn PFFs exhibited significant pHrodo uptake, with a reduction in aSyn-mVenus fluorescence lifetime already apparent on Day 2 in the representative example of Fig. 6(a). In this case, by Day 3, the pHrodo signal had disappeared, and a clear redistribution of the aSyn-mVenus fluorescence was observed in the soma, accompanied by a further decrease in aSyn-mVenus fluorescence lifetime. After 4 days, this particular neuron died. Conversely, neurons treated with mouse aSyn PFFs generally exhibited a strong pHrodo signal from Days 1 through 4, with no indication of aSyn-mVenus redistribution in the soma or a decrease in fluorescence lifetime, as illustrated by the representative case shown in Fig. 6(b). Our results indicate that in the majority of neurons exposed to mouse aSyn PFFs, neither endocytic seed escape nor aSyn-mVenus aggregation occurred over the entire time course. Together, these findings support our earlier interpretation that the escape of fibrillar seeds from the endolysosomal compartment (as indicated by the loss of pHrodo signal) is closely associated with aSyn-mVenus aggregation in neurons.

**Figure 6:**
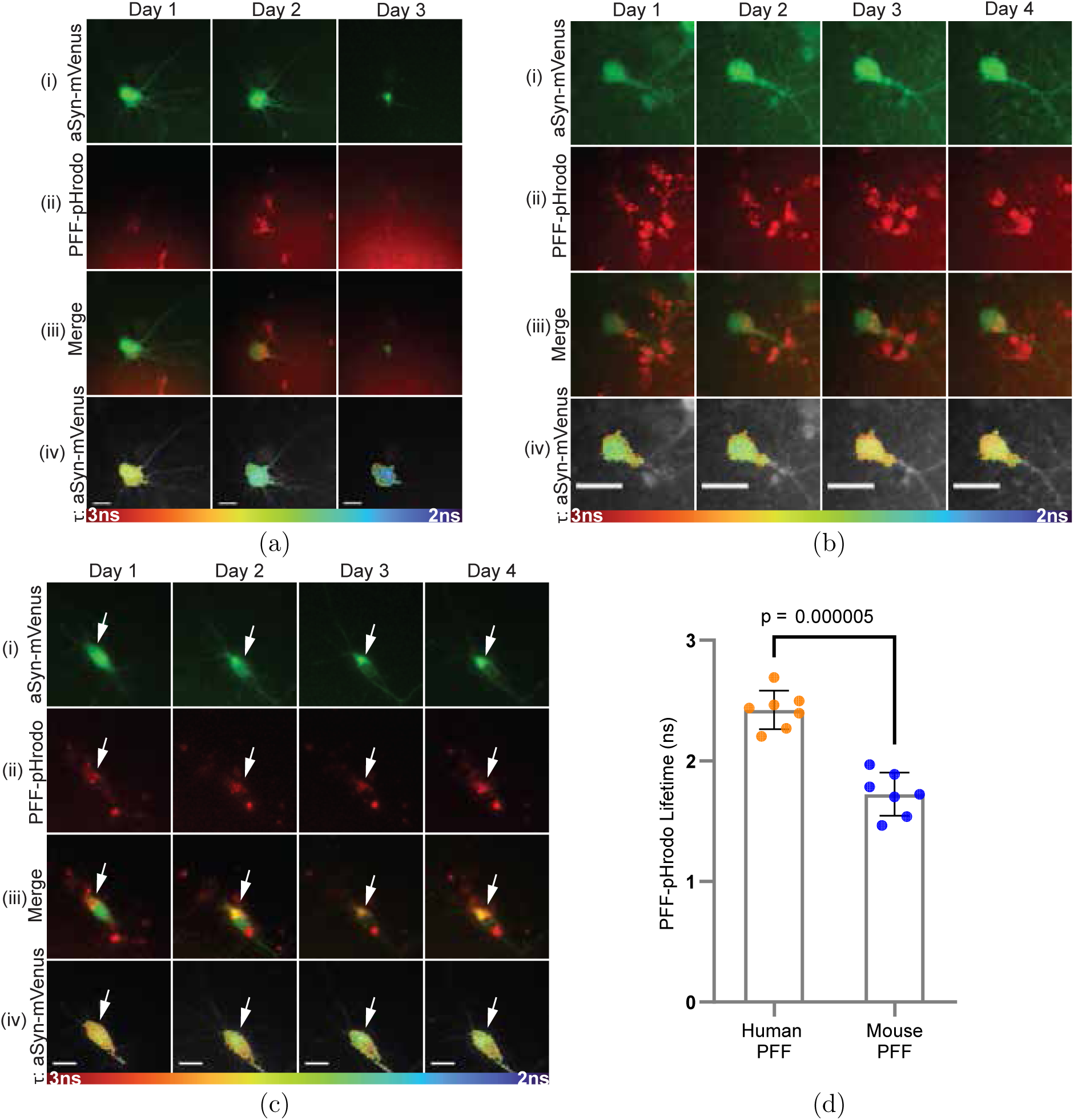
PFF retention is generally (but not always) associated with a lack of seeded aggregation. (a)-(c) Images of representative live cortical neurons transduced with adenovirus encoding aSyn-mVenus and treated with pHrodo-labeled aSyn PFFs, recorded from 3 to 5 days (identified as Day 1 through Day 3) (a) or 3 to 6 days (identified as Day 1 through Day 4) (b, c) post-PFF treatment. The images depict the fluorescence intensity of aSyn-mVenus (i) and pHrodo (ii), the merged intensity signals (iii), and the corresponding aSyn-mVenus lifetime map (iv) of neurons showing evidence of seed escape after treatment with A53T aSyn PFFs (a), or seed retention after treatment with mouse aSyn PFFs (b, c). (a) and (c) show aSyn-mVenus aggregation with the white arrows in (c) highlighting the site of aggregation that also contained PFF-pHrodo inclusions. (b) depicts a lack of aSyn-mVenus aggregation. Scale bars in (a)-(c): 15 *µ*m. (d) Graph showing the aSyn PFF-pHrodo fluorescence lifetime for neurons treated with human A53T aSyn PFFs at the final timepoint before escape or mouse aSyn PFFs on the final day of retention (n = 7 neurons), indicating a significantly lower lifetime for neurons treated with mouse aSyn PFFs. The data are shown as the mean +*/*− SEM (*p <* 0.000005, unpaired t-test).

In a small subset of neurons treated with mouse aSyn PFFs, we observed a retention of punctate pHrodo signal, likely reflecting endolysosomal localization, from Days 1 through 4, coupled with a gradual redistribution of aSyn-mVenus fluorescence that matched the punctate pattern (Fig. 6(c), white arrows) and was accompanied by a progressive decrease in aSyn-mVenus lifetime. This spatial redistribution in fluorescence intensity, together with the reduced lifetime, implies that the cytosolic protein translocated into the endocytic compartment, where it underwent seeded aggregation in association with internalized aSyn PFFs. We validated the findings presented in Fig. 6(c) and reinforced our conclusions by confirming that the fluorescence lifetime of aSyn-mVenus is independent of pH. This was achieved by showing that the lifetime of purified aSyn-mVenus was similar in acidic and neutral en-vironments environments (2.75 ns and 2.80 ns in a low-pH or neutral buffer, respectively; Table 1).

**Table 1:** Fluorescence lifetimes (in ns) of recombinant aSyn variants in solutions of different pH.

| aSyn variant | pH 5.5 | pH 7.4 |
| --- | --- | --- |
| aSyn-pHrodo (monomer) | 2.17 | 2.14 |
| aSyn-pHrodo (fibril) | 1.14 | 1.20 |
| aSyn-mVenus (monomer) | 2.75 | 2.80 |

The evidence presented here that seeded aSyn aggregation can, in some cases, occur within the endolysosomal compartment aligns with the hypothesis that early stages of aSyn self-assembly are favored in the lysosome due to several factors: the high effective concentration of aSyn, the acidic environment, the presence of enzymes capable of modifying internalized seeds, and the potential for aggregation-promoting interactions between aSyn and lysosomal membranes.^32,33^ Overall, our results suggest that endocytic escape, indicated by the loss of pHrodo signal, is associated with aSyn-mVenus aggregation, whereas retention of the pHrodo signal is primarily correlated with the absence of aggregation (in 7 cases of retention, only 2 were associated with aggregation).

Fluorophores such as mVenus, which are derived from green fluorescent protein (GFP), are known to undergo quenching in acidic environments, leading to a reduction in intensity.^34,35^ We note that such a reduction in quantum yield is not necessarily associated with a reduction in lifetime (see Table 1). We verified that aSyn-mVenus is still detectable in acidic environments, albeit with a 2-fold decrease in intensity at acidic compared to neutral pH (Supplementary Fig. 2). To mitigate this decrease in signal, and also to reduce the impact of photobleaching associated with repeatedly exposing the same regions to the laser over multiple days, we increased the gain on the gated intensifier over subsequent days. The resulting signal amplification ensured similar intensity counts across days.

A possible alternative explanation for the decrease in aSyn-mVenus lifetime in the case of PFF-pHrodo retention is that, based on their emission and absorption spectra, a Förster resonance energy transfer (FRET) interaction ^36^ could exist between mVenus and pHrodo, where mVenus acts as the donor. In this case, a lifetime decrease could be attributed to FRET, rather than self-quenching due to molecular crowding indicative of aggregation. However, even if the lifetime decrease were due to FRET, we could still conclude that aSyn-mVenus is interacting with PFF-pHrodo in the endocytic compartment, and this would in turn imply that seeding could occur under these conditions. Discrimination between a FRET or self-quenching effect could be achieved by using a longer wavelength red dye as a PFF label that affords no spectral overlap with mVenus. In such an experiment, a measured decrease in aSyn-mVenus fluorescence lifetime could be solely attributed to self-quenching.

To investigate why human aSyn PFFs exhibited more efficient endocytic escape compared to mouse aSyn PFFs, we analyzed the fluorescence lifetimes of pHrodo-labeled seeds of both types. In initial control experiments, we measured the fluorescence lifetime of pHrodo conjugated to recombinant aSyn monomer or human PFFs in solutions of varying pH to ensure that the pHrodo lifetime could reliably report on the aSyn assembly state without being influenced by pH. pHrodo conjugated to monomeric aSyn showed similar lifetimes in both acidic and neutral buffers (2.17 ns and 2.14 ns, respectively, Table 1), confirming that the pHrodo lifetime is pH-independent under these conditions. Additionally, pHrodo exhibited a significantly shorter lifetime when bound to PFFs compared to monomeric aSyn in acidic solution (1.14 ns versus 2.17 ns, Table 1), indicating that the lifetime effectively reports on the aSyn assembly state. Next, cultures treated with pHrodo-labeled, human aSyn PFFs were examined the day before endocytic escape occurred (as indicated by the loss of pHrodo signal), whereas cultures exposed to pHrdod-labeled, mouse aSyn PFFs were analyzed on the final day of seed retention (i.e., Day 4). The mean fluorescence lifetime of pHrodo associated with human PFFs was significantly greater than that of the dye attached to retained mouse PFFs (Fig. 6(d)), suggesting that the mouse PFFs were larger or denser than their human counterparts. Based on these findings, it can be inferred that smaller, less dense aggregates are more likely to escape from the endolysosomal compartment. These results are consistent with other studies that demonstrated that larger aggregates, for instance those that could be formed from human E46K aSyn, did not lead to endolysosomal vesicle rupture compared to smaller aggregates formed from human WT aSyn.^13^ Additionally, we observed an increase in lifetime of retained mouse PFFs from Day 1 to Day 2, suggesting that aSyn seeds may undergo partial dissociation in acidic compartments (Supplementary Fig. 3).

It remains unclear why some neurons retaining internalized aSyn PFFs exhibited aSyn-mVenus aggregation within the endolysosomal compartment, whereas others did not. One possible explanation is that the translocation of cytosolic aSyn into the endolysosomal compartment (e.g., via chaperone-mediated autophagy ^37^) might be a relatively rare event in the neuronal culture model used here. While we observe this event via a redistribution of aSyn-mVenus intensity signal that matches the geometry of retained PFF-pHrodo, this could be explored more precisely by monitoring the fate of aSyn conjugated to a pH-sensitive FP via confocal microscopy.

### 3.5 aSyn Aggregates Accumulate in Regions of Microtubule Dis-ruption within Neuronal Processes

A final set of experiments was carried out to investigate the origin of the dashed patterns of aSyn-mVenus fluorescence intensity observed in PFF-treated neurons exhibiting signs of seeded aSyn aggregation (for example, in Figs. 3 (b) and (c)). It has been shown that aSyn interacts with tubulin in microtubules^38^ and can influence microtubule polymerization.^39^ In addition, microtubule transport has been proposed as a means for aggregate deposition along axons,^40^ and disruptions of the microtubule cytoskeletal structure are commonly observed in neurodegenerative diseases. ^41^ Based on this evidence, we hypothesized that the dashed pattern could result from the deposition of aSyn-mVenus protein along microtubule tracts within neuronal processes, potentially driven by a disruption of the microtubule structure.

To address the question of aSyn deposition along microtubule tracts, rat cortical neurons expressing aSyn-mVenus were incubated in the absence or presence of human A53T aSyn PFFs for 6 days, fixed, and stained for beta-tubulin with Alexa fluor 594. Our results are shown in Fig. 7. In neurons cultured without PFFs, the aSyn-mVenus signal was distributed throughout the neuron and extensively overlapped with beta-tubulin immunoreactivity, except in a distinct hollow region at the center of the beta-tubulin signal, corresponding to the nucleus (Fig. 7, top panel). In contrast, in PFF-treated neurons, aSyn-mVenus fluorescence redistributed to a perinuclear region within the soma, coinciding with a depletion of the beta-tubulin signal (Fig. 7, middle panel). A similar pattern was observed in neuronal processes, where beta-tubulin immunoreactivity was absent from ‘dashed’ regions characterized by the accumulation of aSyn-mVenus aggregates (Fig. 7, bottom panel). These results suggest that aSyn aggregates disrupt the microtubule network in both the soma and neuronal processes. This is in agreement with previous reports that aSyn oligomers or fibrils affect microtubule stability, leading to a disruption of the microtubule network.^42^

**Figure 7:**
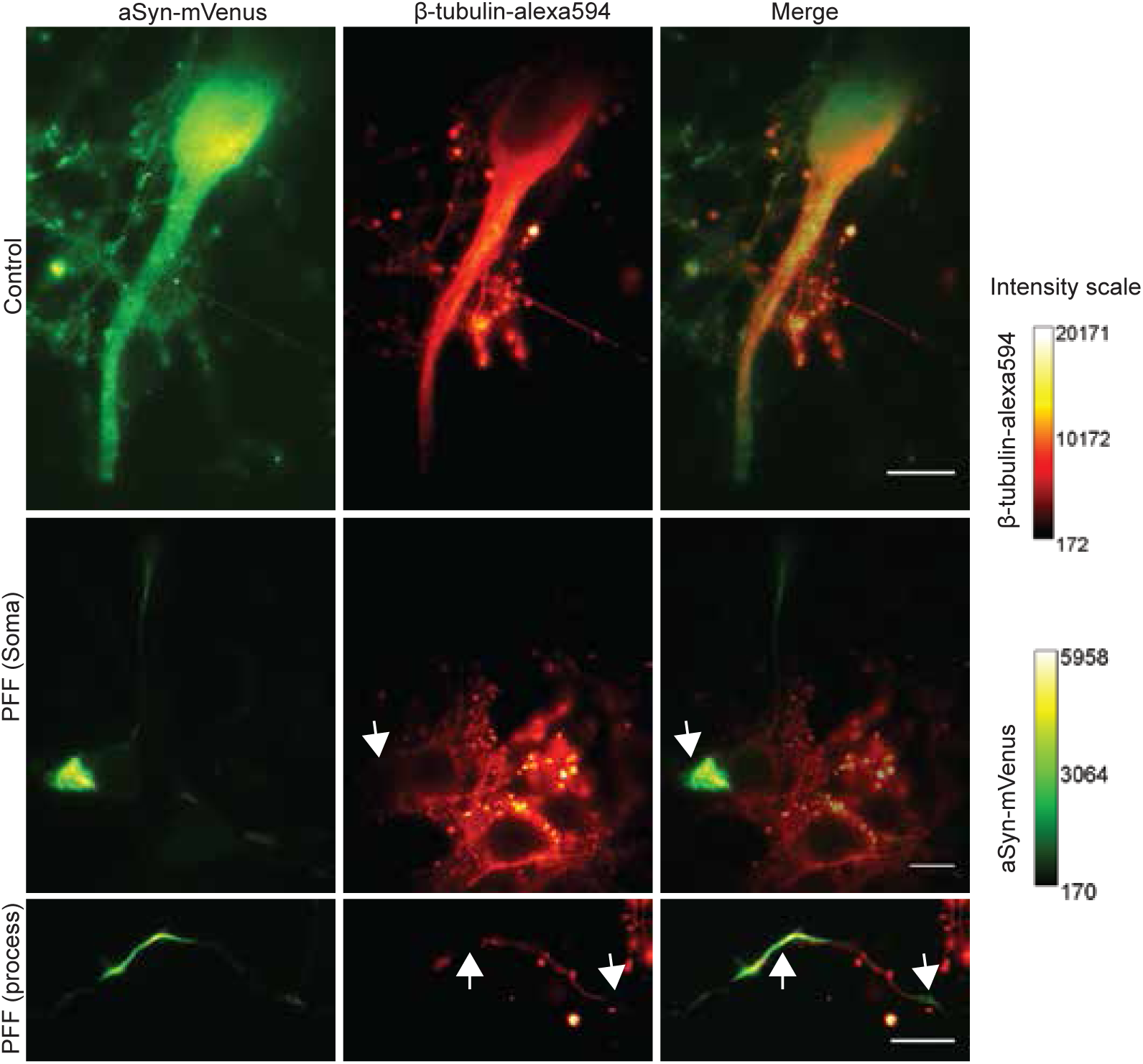
aSyn aggregates disrupt the microtubule network in neurons treated with aSyn PFFs. Rat cortical neurons expressing aSyn-mVenus were incubated in the absence (‘control’, top row) or presence (middle and bottom rows) of human A53T aSyn PFFs for 6 days. The cells were fixed 6 days post-PFF treatment, stained with a primary antibody specific for *β*-tubulin and an Alexa Fluor 594-conjugated secondary antibody, and imaged. The panels depict the fluorescence intensity of aSyn-mVenus (left column) and Alexa Fluor 594 (middle column), alongside the merged intensity signals (right column). White arrows indicate regions in the soma (middle row) or processes (bottom row) of representative PFF-treated neurons with apparent aSyn-mVenus aggregates, accompanied by a corresponding loss of *β*-tubulin-Alexa 594 signal. Scale bar: 10 *µ*m.

Based on these findings, we propose that aSyn aggregates are transported along microtubule tracts, likely as part of a cellular response involving their sequestration into aggre-somes via retrograde transport.^43^ However, during this process, the aggregates destabilize the microtubule network responsible for their transport, leading to their deposition at sites of microtubule disruption. The mechanism underlying this microtubule destabilization remains unclear. The aggregates could either actively depolymerize microtubules through a mechanical interaction, or they could interfere with beta-tubulin polymerization.

## 4 Conclusions

We implemented an advanced imaging platform to track self-quenching-induced fluorescence lifetime changes, providing a sensitive measure of seeded aSyn aggregation in primary rat cortical neurons. This method enabled direct visualization of the seeding mechanism. In contrast, previous efforts to study this phenomenon were limited, as they relied on examining fluorescence intensity puncta to monitor aggregation, an approach that is less sensitive than lifetime measurements. In addition, previous methods lacked sensitivity to fibril location based on pH. Employing our imaging methods in primary neuron cultures enabled us to probe aggregation-induced morphologies uniquely distributed along neuronal processes. By co-staining fixed cells for beta-tubulin, we were able to show that the formation of these structures interferes with microtubule structure.

Our results indicate that endocytic escape and retention of fibrillar seeds can both act as seeding pathways for aggregation (summarized in Fig. 8), although the escape mechanism was predominantly associated with aggregation. This was demonstrated by examining neurons for a decrease in aSyn-mVenus lifetime as a measure of aggregation, along with the presence/absence of a pHrodo signal, which determined the retention/escape of fibrillar seeds (PFFs). There are still open questions regarding the fate of retained fibrils. For instance, in which part of the endolysosomal compartment can seeded aSyn aggregation occur, and what ultimately happens to the aggregate that is formed? One possibility is that the aggregate may exit the endolysosomal compartment, potentially via autophagic or lysosomal secretion,^44,45^ enabling internalization by a neighboring neuron (Fig. 8). Collectively, our findings provide novel insights into the molecular mechanisms driving aSyn pathology propagation and the resulting neuronal dysfunction in synucleinopathy disorders, setting the stage for developing new therapeutic strategies.

**Figure 8:**
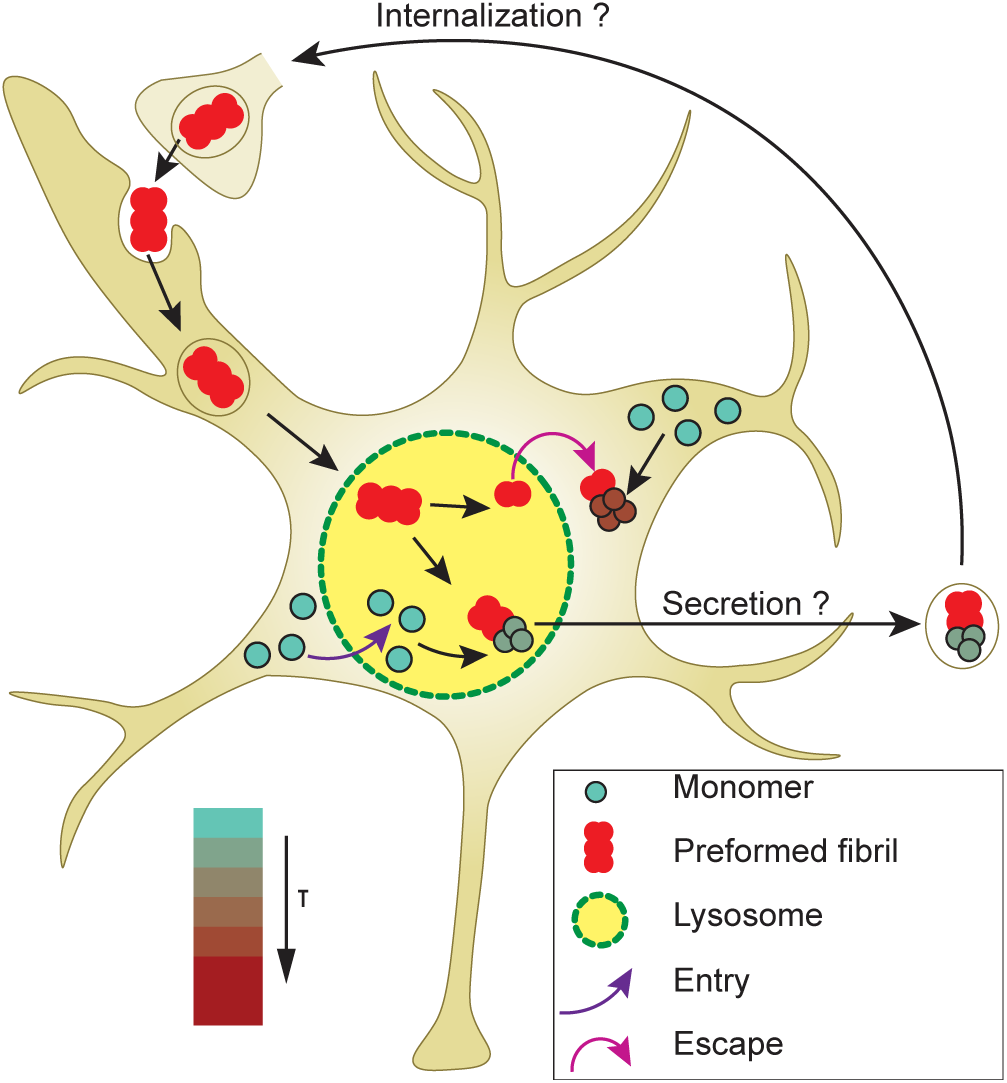
Model illustrating the major findings of this study. aSyn PFFs (shown as a red multi-subunit assembly at the top) are internalized via endocytosis and trafficked to the lysosome. Within the lysosome, PFFs undergo partial dissociation, represented by the fragmentation of the red assembly. Smaller PFF fragments exit the lysosome (indicated by a purple arrow denoting endocytic escape) and recruit cytosolic aSyn monomers (depicted as blue spheres). This interaction promotes seeded aSyn aggregation and, in aSyn-mVenus-expressing cells, results in a decrease in fluorescence lifetime, visualized as a red-shifted gradient. Conversely, larger PFF fragments are retained in the lysosome but can recruit monomeric aSyn transported into this compartment (potentially via chaperone-mediated autophagy). This recruitment leads to seeded aSyn aggregation in the lysosome and, in aSyn-mVenus-expressing cells, a corresponding decrease in aSyn-mVenus lifetime within this compartment. aSyn aggregates formed in the endolysosomal compartment may be released from the neuron, potentially via autophagic or lysosomal secretion. In turn, the secreted aggregates could be internalized by neighboring neurons, contributing to the propagation of pathology.

## 5 Methods

### 5.1 Design of the FLIM Platform

A custom time-domain FLIM setup was constructed around an Olympus iX73 inverted microscope platform. The excitation source was a 1-MHz 1040-nm pump laser coupled with an optical parametric amplifier (Spectra-Physics) to support a broader range of excitation wavelengths (320-1040 nm). The laser was coupled to the microscope with a series of lenses and mirrors. The system was equipped with two image-capturing devices, an sCMOS camera (Photometrics PRIME) capable of up to 100 fps collection rates for intensity-based imaging, and a high-rate image-intensified (PicoStar HR12, LaVision) CMOS (Imager-M-Lite, LaVision) integrated system for fluorescence lifetime recording. A picosecond delay unit (PSD) was used to introduce delays between the intensifier gate and the exciting laser pulse. The delay unit used the electrical trigger output of the laser. For the FLIM measurements, an intensifier gate width of 200 ps was applied and delayed relative to the laser pulse in 200 ps increments to sample the fluorescence decay curve. A pixel-wise mono-exponential fit with the deconvolved temporal gate function was then performed on the data to determine the lifetime.

### 5.2 Expression and Purification of Recombinant aSyn Proteins

Recombinant aSyn variants were purified following a protocol adapted from Zhang *et al.* (2013)^46^ with modifications described by Volpicelli *et al.* (2014).^47^ *E. coli* BL21 (DE3) cells were transformed with the bacterial expression vector pT7-7 encoding A53T aSyn, mouse aSyn, or human wild-type (WT) aSyn-mVenus. A single colony was used to inoculate LB medium supplemented with ampicillin (100 *µ*g/L), and the culture was incubated at 37*^◦^*C until the optical density at 600 nm (OD600) reached 0.5-0.6. Protein expression was induced with isopropyl *β*-D-1-thiogalactopyranoside at a final concentration of 1 mM, followed by incubation at 37*^◦^*C for 4 h. The cells were then harvested by centrifugation.

Cells expressing untagged aSyn (A53T or mouse aSyn) were resuspended in lysis buffer A (10 mM Tris HCl, pH 8.0, 1 mM EDTA, 0.25 mg/mL lysozyme) and lysed using a French press cell disruptor (Thermo Electron, Waltham, MA) at *>* 1000 psi. The lysate was treated with 0.1% (w/v) streptomycin sulfate to precipitate DNA and then clarified by centrifugation. The supernatant was subjected to partial purification via two successive ammonium sulfate precipitations (30% and 50% saturation) at 4*^◦^*C. The resulting pellet was resuspended in 10 mM Tris-HCl (pH 7.4), and the suspension was heated to 100*^◦^*C for 15 min. Following centrifugation at 13,500 × g at 4*^◦^*C for 20 min to precipitate denatured proteins, the supernatant was filtered through a 0.22-*µ*m membrane. aSyn was purified via successive fractionations using: (i) a HiLoad 16/600 Superdex 200 pg size exclusion column (Cytiva, Marlborough, MA), with elution performed in 10 mM Tris HCl (pH 7.4); and (ii) a HiPrep Q HP 16/10 anion-exchange column (Cytiva), with elution performed using a linear gradient of 25 mM to 1 M NaCl in a buffer consisting of 10 mM Tris HCl (pH 7.4) and 1 mM EDTA. Cells expressing aSyn-mVenus were resuspended in lysis buffer B (50 mM Tris-HCl, pH 8.0, 50 mM NaCl, and 5 mM imidazole) supplemented with 0.2 mg/mL lysozyme and 1 mM PMSF. The cells were lysed on ice using a probe sonicator (Fisherbrand Model 120) over 12 cycles, each consisting of a 15-s pulse followed by a 45-s recovery period. The lysate was filtered through a 0.22-*µ*m membrane, and aSyn-mVenus was purified via successive fractionations using: (i) a 5 mL HisTrap HP column (Cytiva), with elution performed using a linear gradient of 5 to 500 mM imidazole in a buffer consisting of 50 mM Tris-HCl (pH 8.0) and 50 mM NaCl; and (ii) a HiLoad 16/600 Superdex 200 size exclusion column, with elution performed in a buffer consisting of 25 mM Tris HCl (pH 7.4), 100 mM NaCl, and 0.01% NaN_3_.

Fractions enriched with aSyn or aSyn-mVenus (identified via SDS-PAGE with Coomassie blue staining) were pooled, and the solution was dialyzed against PBS (10 mM phosphate buffer, 2.7 mM KCl, and 137 mM NaCl, pH 7.4) in the case of untagged aSyn, or left undialyzed in the case of aSyn-mVenus. The purified protein was stored at-80*^◦^*C until use, with a final purity of approximately 95%.

### 5.3 Preparation of aSyn Preformed Fibrils

Purified human A53T aSyn or mouse aSyn was concentrated to 5 mg/mL (347 *µ*M) using a 10 kDa molecular weight cutoff (MWCO) spin filter (Sartorius, Columbus, OH) operated at 4,500 × g. The concentrated protein solution (0.5 mL) was filtered through a 0.22-*µ*m syringe filter (Fisherbrand polyethersulfone, 33 mm) and incubated in a sterile 1.5-mL mi-crocentrifuge tube at 37*^◦^*C for 7 days with continuous shaking at 123 × g in a Thermomixer (BT LabSystems BT917). The resulting fibrils were collected by centrifugation at 13,000 × g for 15 min and resuspended in 250 *µ*L of Dulbecco’s phosphate-buffered saline (DPBS; Cytiva). An aliquot of the fibril suspension was treated with guanidine hydrochloride (final concentration, 8 M) and incubated at 22*^◦^*C for 1 h to dissociate fibrils into monomers. The aSyn concentration was measured via absorbance at 280 nm using a Nanodrop spectrophotometer, with extinction coefficients of 5960 M^-1^cm^-1^ and 7450 M^-1^cm^-1^ for human and mouse aSyn, respectively. Fibrils were resuspended to a final concentration of 5 mg/mL, divided into 257 *µ*L aliquots, and stored at-80*^◦^*C. Before use, fibril suspensions were sonicated in ethanol-sterilized tubes (Active Motif) using a cup horn sonicator (Qsonica Q700) set to 30% power (100 W/s) with a cycle of 3 s on and 2 s off, for a total ‘on’ time of 8 min. Fibrils prepared from mouse or human aSyn underwent a total of 2 or 3 rounds of sonication, respectively. The bath temperature was maintained between 5*^◦^*C and 15*^◦^*C throughout the sonication.

### 5.4 Labeling of Preformed Fibrils with pHrodo Red

Fibrillar aSyn was labeled with an N-hydroxysuccinimide (NHS) ester derivative of the pH-sensitive dye pHrodo Red (Thermo Fisher Scientific). Unsonicated fibrils, prepared from recombinant A53T aSyn or mouse aSyn, were buffer-exchanged into PBS (pH 7.4) using 10 kDa MWCO filters (Thermo Fisher Scientific). The fibrils were then incubated with a 1-to 5-fold molar excess of dye (relative to the monomer equivalent of fibrillar aSyn) at 22*^◦^*C for 1 h in a light-protected tube. Stock dye solutions of 10 to 50 mM were used to ensure that the final DMSO concentration in the protein-dye mixture was ≤ 5% (v/v). Unreacted dye was removed by centrifugation using a Zeba spin desalting column (Thermo Fisher Scientific).

### 5.5 Fluorescence Lifetime Analysis of Recombinant aSyn Variants

Monomeric and fibrillar A53T aSyn conjugated with pHrodo Red, along with recombinant A53T aSyn-mVenus, were diluted to a final concentration of 1 *µ*M in PBS (pH 7.4) or 10 mM MES (pH 5.5). For fluorescence lifetime measurements, a 20 *µ*L aliquot of each solution was placed on a glass slide and sealed with a coverslip to prevent evaporation. Imaging was performed at 40× magnification under appropriate excitation and emission settings for each fluorophore.

### 5.6 Preparation of aSyn A53T-mVenus Adenovirus

An adenoviral construct encoding human aSyn A53T-mVenus was produced using the Vi-rapower Adenoviral Expression System from Invitrogen (Carlsbad, CA). The aSyn A53T-mVenus cDNA was subcloned as a KpnI-XhoI fragment into a variant of the entry vector pENTR1A carrying the human synapsin promoter (h-syn-P).^48^ Phusion High-Fidelity DNA polymerase or Q5 Hot Start High-Fidelity DNA polymerase (NEB, Ipswich, MA) was used to carry out the PCR. The sequences of the oligonucleotide primers used to generate the construct are listed in the supplementary material (Supplementary Table 1). The ligation reaction was carried out using the NEBuilder HiFi DNA Assembly kit (NEB). The insert from the pENTR1A construct was transferred into the ‘promoter-less’ pAd/PL-DEST5 adenoviral expression vector^48^ via Gateway recombination cloning (Invitrogen). The DNA sequence of the insert was verified by Sanger sequencing (Genewiz, South Plainfield, NJ). The resulting adenoviral construct was packaged into virus via lipid-mediated transient transfection of the HEK 293A packaging cell line. Adenoviral titers were determined using the Adeno-X qPCR titration kit from Takara Bio USA (Mountain View, CA).

### 5.7 Preparation and Treatment of Primary Cortical Cultures

Primary cortical cultures were prepared as described^47^ by dissecting day 17 embryos obtained from pregnant Sprague–Dawley rats (Envigo, Indianapolis, IN) using methods approved by the Purdue Animal Care and Use Committee. Briefly, the cortical layers were isolated stereoscopically and dissociated by incubation with papain (20 U/mL) in sterile Hank’s Balanced Salt Solution (HBSS) at 37*^◦^*C for 45 min. The dissociated cells were plated on poly-D-lysine-coated 8-well chambered glass plates (Cellvis) at a density of 75,000 cells/well in Neurobasal media supplemented with 2% (v/v) B-27 supplement, 5% (v/v) FBS, 1% (v/v) GlutaMAX, 50 U/mL penicillin, and 50 *µ*g/mL streptomycin. The next day, the plating media was replaced with Neurobasal media plus 2% (v/v) B-27 supplement, 1% (v/v) GlutaMAX, 10 U/mL penicillin, and 10 *µ*g/mL streptomycin. After 6 days in vitro (DIV = 6), the cultures were transduced with adenovirus encoding aSyn-mVenus under the control of the synapsin promoter at a multiplicity of infection (MOI) of 5. The cultures were treated 24 h later with aSyn PFFs (final concentration, 6 *µ*g/mL) and incubated for 6 days for imaging following fixation, or 6 to 7 days for live-cell imaging experiments. Control cultures were transduced with aSyn-mVenus virus and incubated without PFFs.

### 5.8 FLIM Analysis of Fixed Neurons

Cells were fixed 6 days post-PFF treatment with 4% (w/v) paraformaldehyde (PFA) in PBS in the absence or presence of 1% (v/v) Triton X-100 for 15 min. mVenus fluorescence lifetime data were recorded from 10 fields of view per well of the 8-well chambered glass plate at a magnification of 40 X. aSyn-mVenus was excited at 488 nm, with emission collected through a 530/40 nm band-pass filter.

### 5.9 FLIM Analysis of Live Neurons

Live-cell imaging was conducted from 3 to 6 or 7 days post-PFF treatment using a custom-built FLIM system. During imaging, the cells were maintained on the microscope stage (encoded MCL-MOTNZ, Mad City Labs) inside a live-cell chamber (Okolabs) at 37*^◦^*C and 5% CO_2_. On the first day of imaging (3 days post-PFF treatment), 20 fields of view per well were selected from the 8-well chambered glass plate based on healthy neuron morphology and an optimal signal-to-noise ratio, with stage positions saved for subsequent imaging. Each field-of-view was imaged at a magnification of 20 X. Fluorescence lifetime data were recorded from the same fields on the first day and all subsequent imaging days. aSyn-mVenus was excited at 488 nm, with emission collected through a 530/40 nm band-pass filter. pHrodo Red was excited at 550 nm, with emission collected through a 600/50 nm band-pass filter.

### 5.10 Immunocytochemical Analysis

Cells expressing aSyn-mVenus were fixed 6 days post-PFF treatment with 4% (w/v) PFA in PBS (pH 7.4) (without 1% (v/v) Triton X-100) for 15 min. Following fixation, the cells were incubated in the dark at 22*^◦^*C for 1 h in blocking buffer (PBS, pH 7.4), 1% (w/v) BSA, 10% (v/v) FBS, and 0.3% (v/v) Triton X-100). The cells were then incubated with rabbit *β*3-tubulin primary antibody (D65A4, Cell Signaling Technology, Danvers, MA), diluted 1:500 in dilution buffer (PBS (pH 7.4), 1% (w/v) BSA), at 4*^◦^*C for 24 h. After washing with PBS (pH 7.4), the cells were treated with an Alexa Fluor 594-conjugated goat anti-rabbit secondary antibody, diluted 1:1000 in dilution buffer, at 22*^◦^*C for 1 h. After a final wash with PBS (pH 7.4), the samples were sealed with parafilm for imaging.

### 5.11 Statistical Analysis

The data were analyzed with an unpaired t-test (for experiments with 2 groups) using Graph-Pad Prism version 8.0 (La Jolla, CA).

## Acknowledgement

This work was supported in part by the National Science Foundation (CBET 1937986 and EAGER 2330643), the National Institutes of Health (R21 NS105048 and NS135424), and by the Michael J. Fox Foundation.

## Supporting Information Available

An associated supplementary file contains the following referenced figures: the MATLAB generated live-cell imaging GUI used to analyze data, a comparison of aSyn-mVenus fluorescence intensity in acidic (MES) and neutral (PBS) buffer conditions, and a graph of retained mouse PFF-pHrodo lifetimes tracked over several days. Additionally, the oligonucleotide sequences used to generate the aSyn A53T-mVenus adenoviral construct are presented in Supplementary Table 1.

