## Supplementary Material for "Mechanisms of Alpha-Synuclein Seeded Aggregation in Neurons Revealed by Fluorescence Lifetime Imaging"

This document contains the MATLAB generated live-cell imaging GUI used to analyze data (Fig. 1), a comparison of aSyn-mVenus fluorescence intensity in acidic (MES) and neutral (PBS) buffer conditions (Fig. 2), and a graph of retained mouse PFF-pHrodo lifetimes tracked over several days (Fig. 3). Additionally, the oligonucleotide sequences used to generate the aSyn A53T-mVenus adenoviral construct are presented in Supplementary Table 1.

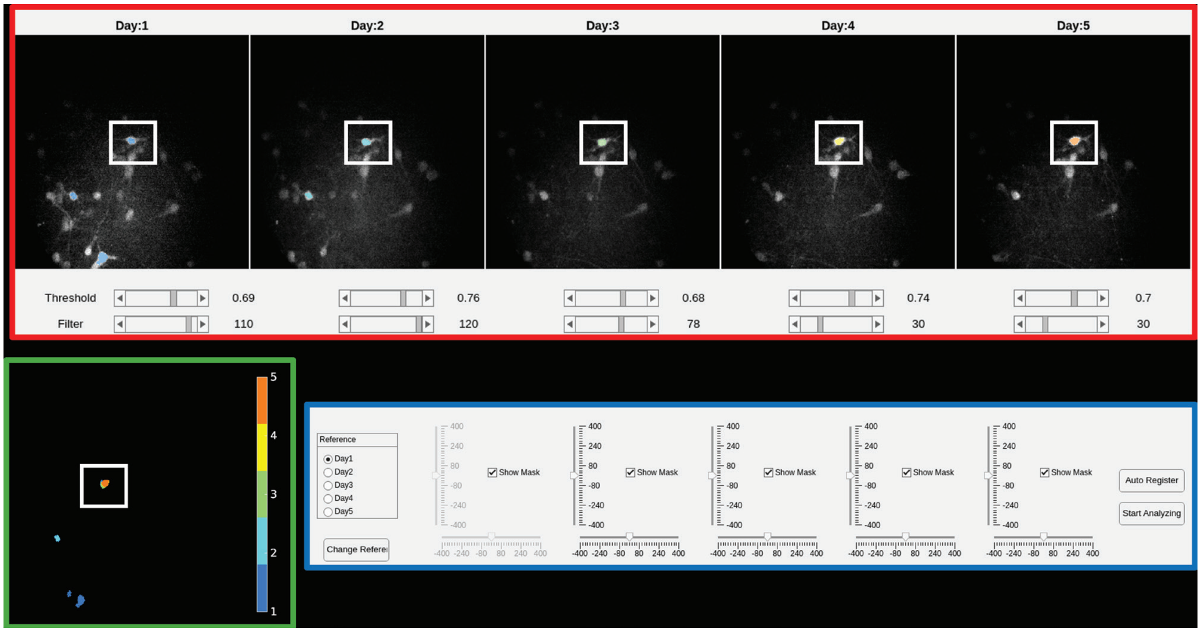


Figure 1: Live-cell imaging GUI. The red panel shows intensity images of the same region of a well imaged over 4 days. For each day, the white square highlights a neuron that is extracted by applying a threshold controlled separately for each image. The green panel shows the extracted threshold masks from each intensity image. The bar on the right shows the mask colors, with each color representing each day of imaging. The white square shows that the masks overlap for the 4 days for the neuron highlighted in the red panel. The blue panel shows control buttons for each image where the images' lateral and vertical positions can be adjusted relative to a selected reference image to ensure registration of the thresholded masks for the neuron being extracted. The thresholded mask is then applied to each image, and the lifetime in the thresholded region is measured on a pixel-wise basis to produce a lifetime map showing the lifetime of each pixel in the map.

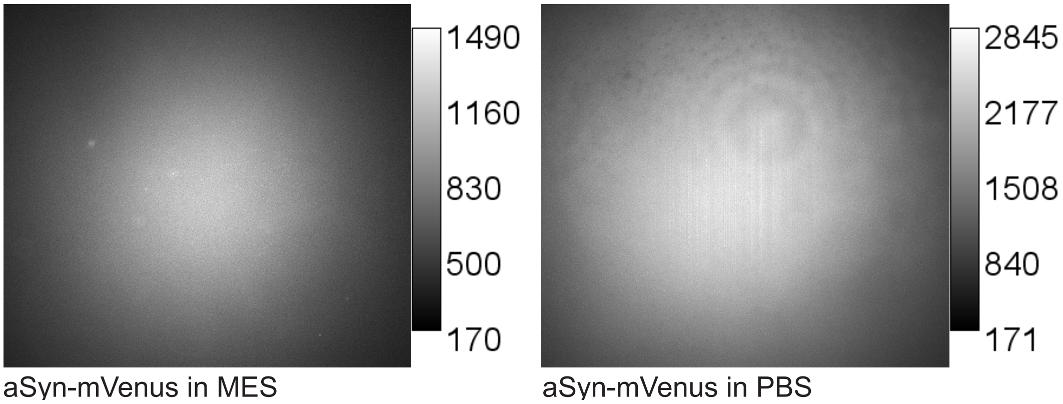


Figure 2: Intensity images of purified aSyn-mVenus in MES and PBS buffers demonstrating that aSyn-mVenus has a 2x lower yield in an acidic environment (MES) compared to a neutral one (PBS).

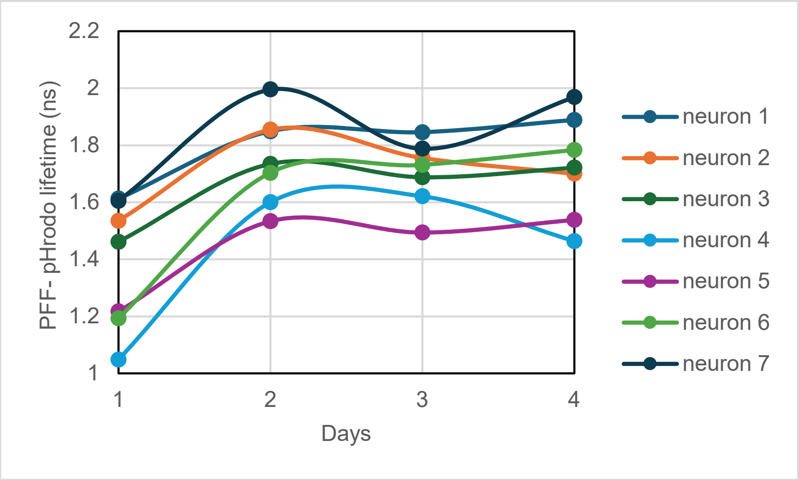


Figure 3: Graph showing the average lifetime of retained mouse PFF-pHrodo over 4 days for 7 different neurons. The average PFF-pHrodo lifetime for each neuron was calculated from a distribution of pixels within the neuron with PFF-pHrodo signal. The average lifetime was measured over four days from the same distribution of pixels determined from a thresholded mask (described in Fig. 1). The overall trend observed is that the mouse PFF-pHrodo lifetime increased from Day 1 to Day 2 and then plateaued over the remaining days.

Table 1: Sequences of the oligonucleotide primers used to generate the aSyn A53T-mVenus adenoviral construct.

| Primer | Sequence (5'-3') |
|---|---|
| aSyn-linker-mVenus-For | GACTACGAACCTGAAGCCGCTCCAGTAGCTACAATGGTGAGCAAGGGCG |
| mVenus-XhoI-pENTR-Rev | GCTGGGTCTAGATATCTCGAGTTACTTGTACAGCTCGTCCATG |
| pENTR-hsyn-KpnI-For | CAAGGATCCACCGGTACCGCCGCCAC |
| aSynA53T-Rev | CCACTGTTGTCACACCATGCACCACTCC |
| aSynA53T-For | GGTGTGACAACAGTGGCTGAGAAGACCAAAG |
| aSyn-EcoRI-Rev | CCACAGGCATATCTTCCAGAATTC |
